# Probing the substrate binding-induced conformational change of a ZIP metal transporter using a sandwich ELISA

**DOI:** 10.1101/2025.03.09.642161

**Authors:** Yao Zhang, Ryan Hu, Min Su, Jian Hu

**Affiliations:** Department of Biochemistry and Molecular Biology, Michigan State University, East Lansing, MI 48824; Electron Microscopy Core, University of Missouri, MO 65211; Department of Biochemistry, University of Missouri, MO 65211; Department of Chemistry, Michigan State University, East Lansing, MI 48824

**Keywords:** ZIP, zinc, cysteine accessibility, ELISA, conformation, transporter

## Abstract

Zrt-/Irt-like proteins (ZIPs), a family of divalent metal transporters, are crucial for maintaining the homeostasis of zinc, an essential trace element involved in numerous biological processes. While extensive research on the prototypical ZIP from *Bordetella bronchiseptica* (BbZIP) have suggested an elevator transport mechanism, the dynamic conformational changes during the transport cycle have not been thoroughly studied. In this work, we developed a sandwich ELISA using a custom anti-BbZIP monoclonal antibody to investigate the conformational change induced by the metal binding to the transport site. This was achieved by determining the accessibility of a cysteine residue introduced at a position exposed to the solvent only when the transporter adopts an outward-facing conformation. This assay allowed us to report the dissociation constants of BbZIP for Zn^2+^ and Cd^2+^ at low and sub-micromolar levels, respectively. Notably, the installation of a positive charge at the M2 site drastically reduced metal binding at the M1 site, consistent with an auxiliary role for the M2 site in metal transport. We also demonstrated that this assay can be used to rapidly screen variants for subsequent structural study. We anticipate that other transporters where substrate binding induces large conformational changes can also be studied using this method.

## Introduction

Zinc is an essential trace element in living organisms [1], as it is extensively involved in enzyme catalysis, protein structure stabilization and gene expression regulation [2-5]. Among the zinc transporters that play a central role in maintaining zinc homeostasis [6], Zrt-/Irt-like proteins (ZIPs) form a large solute carrier protein family (SLC39A) that is ubiquitously expressed throughout the kingdom of life [7-10]. Zn^2+^ is the primary substrate for most characterized ZIPs, but other divalent *d*-block metals can also be transported by some family members. For instance, human ZIP8 and ZIP14 are known to be more promiscuous than other human ZIPs, transporting Zn^2+^, Mn^2+^, Fe^2+^ and the toxic Cd^2+^ [11-15]. IRT1, a root-expressed plant ZIP, is essential for Fe^2+^ uptake, but also exhibits activities for Zn^2+^, Mn^2+^ and Cd^2+^, and is responsible for cadmium uptake from soil under iron-deficient conditions [16-18]. Acting as importers, bacterial ZIPs at the plasma membrane mediate metal uptake from the environment [19] and are part of the metal uptake system of bacterial pathogens to escape the metal deprivation imposed by the host during infection [20]. In eukaryotic cells, ZIPs are widely distributed in the endomembrane system – plasma membrane ZIPs transport metals into the cytoplasm, whereas those on the membrane of intracellular organelles/vesicles are responsible for metal release from storage [21, 22]. Due to their broad and critical functions in many physio/pathological events, some human ZIPs have been identified as drug targets [21, 23-32], warranting fundamental biochemical and mechanistic studies.

The last decade witnessed the rapid progress in the structural biology study of a bacterial ZIP from *Bordetella bronchiseptica* (BbZIP) [33]. Structures solved by x-ray crystallography and cryo-EM have revealed a novel transporter fold consisting of two structurally independent domains [34-38]. Evolutionary covariance analysis, symmetry-based modeling, computational simulations, and biochemical studies have consistently supported an elevator transport mode of this prototypic ZIP [35-37]. The unique hinge motion of the transport domain during conformational change, which were revealed in both experimentally solved structures and metadynamic simulations [35], distinguishes BbZIP from other known elevator transporters. Three specific metal binding sites along the transport pathway have been identified in various BbZIP structures, including a binuclear metal center (M1 and M2) at the transport site and an inhibitory metal binding site (M3) at the exit of metal release pathway [34, 35, 38]. Sequence analysis have shown that M1 is the most conserved metal binding across the entire ZIP family, whereas M2 is present in many but not all ZIPs [39]. The M3 site, which is located in the cytosolic loop between TM3 and TM4, is the most variable metal binding site [9, 35, 38]. Functional comparison between M1 and M2 on human ZIP4 indicated that M1 is essential for zinc transport whereas M2 may play an accessory role as M2 deletion only modestly reduced the transport rate, but the exact function of M2 still remains to be clarified. The apparent *K*_M_ values of many ZIPs have been measured in kinetic studies performed on live cells, allowing the binding affinity of the transport site to be estimated for various metal substrates [17, 40-44]. However, biochemically characterizing the metal binding to BbZIP is challenging because the purified BbZIP in the apo state is unstable in detergent micelles. As a result, although BbZIP, as a prototypical member of the ZIP family, have been extensively studied for years, there is still no report about the binding affinity of BbZIP for its likely physiological substrate Zn^2+^ and a surrogate substrate Cd^2+^.

In this work, based on the elevator transport mode and our previously established cysteine accessibility assay [35-37], we developed a sandwich ELISA to probe the metal induced conformational change of BbZIP in the native membrane. Using this approach, we demonstrated the selective metal binding to BbZIP, measured the binding affinities for Zn^2+^ and Cd^2+^, and studied the effects of mutagenesis on zinc binding and the conformational state of the transporter. These results complement the previous structural and biochemical studies and provide additional insights into the mechanism of metal transport by BbZIP.

## Results and Discussion

### An introduced cysteine residue as a conformation indicator

It has been shown that L200, which is located in the transport domain and being one of the residues that block the pathway toward the periplasm [10], is buried when BbZIP is in the inward-facing conformation (IFC) but exposed to solvent in the proposed outward-facing conformation (OFC) (**Figure 1A**) [35, 36]. Taking advantage of the absence of cysteine residue in wild-type BbZIP, a cysteine was introduced to replace L200 and the accessibility of C200 to thiol reacting agents in the L200C variant can be used to deduce the conformational state of the transporter [35]. In this assay, the exposed C200 will be covalently modified by a low molecular weight thiol reacting agent N-ethylmaleimide (NEM) in the native state, and thus will no longer be able to be labeled by a high molecular weight thiol reacting agent (∼5000 Da), monofunctional PEG-Maleimide 5K (mPEG5K), under denatured conditions. In contrast, C200 buried in protein core cannot be labeled by NEM under native conditions but will be subsequently labeled by mPEG5K upon protein denaturation in SDS and urea. By analyzing of the ratio of the NEM-labeled and mPEG5K-labeled species, which can be estimated in Western blot where the two species can be differentiated by size, one can deduce the transporter’s conformational state. Importantly, since elimination of the M1 metal binding site, which is the most conserved metal binding site and absolutely required for transport [39], rather than the neighboring and more variable M2 site, completely abolished the response to the addition of metal substrates [35], the change of the cysteine accessibility of the L200C variant can therefore be attributed to the conformational change triggered by the metal binding to the M1 site. As reported in our recent study, the addition of the known metal substrates of BbZIP, Zn^2+^ and Cd^2+^, to the membrane fraction of cells expressing the L200C variant resulted in a significant increase of the mPEG5K-labeled species (**Figure 1B**), indicating that the binding of metal substrates to the M1 site drastically reduced the accessibility of C200 due to the substrate binding induced OFC-to-IFC switch. Although this well-established approach can be used to characterize metal binding to the M1 site, one may suffer from the relatively poor accuracy with large variations that are intrinsically associated with the Western blot experiments. To address this issue, using a custom anti-BbZIP monoclonal antibody, which has been validated in our early studies [35, 36], as the coating antibody, we converted this cysteine accessibility assay conducted in test tubes into a sandwich enzyme-linked immunosorbent assay (ELISA) conducted in 96-well plates, allowing us to rapidly and more accurately study the binding of metal substrates to BbZIP that is embedded in the native membrane.

**Figure 1.**
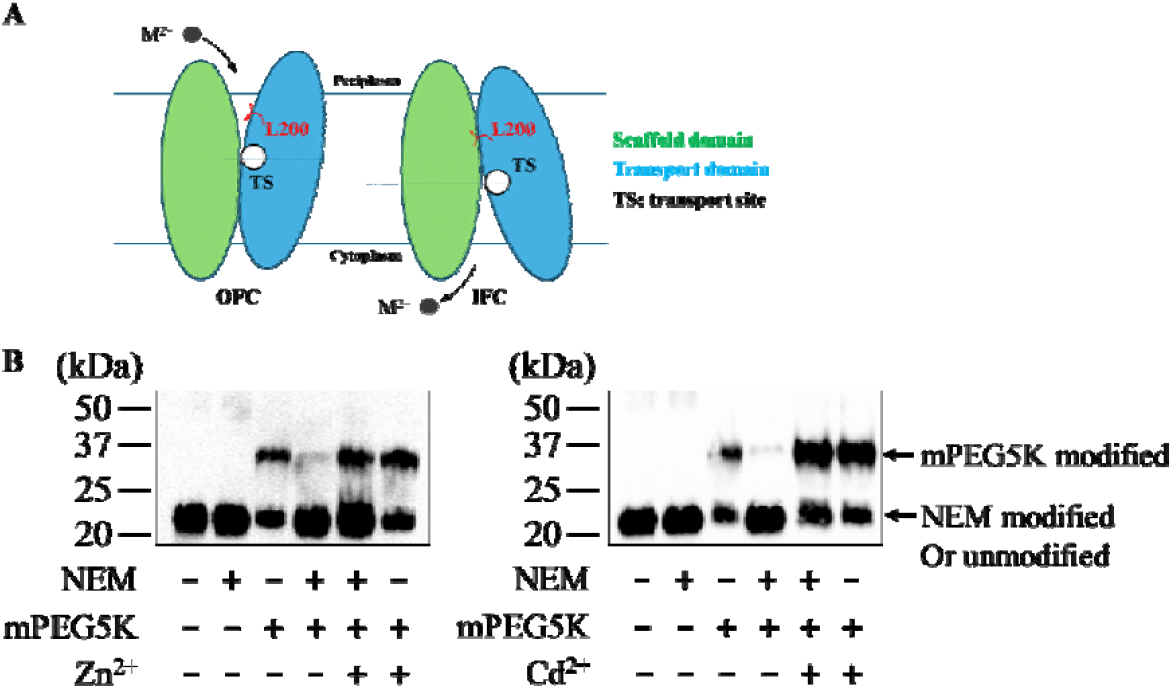
Altered accessibility of L200 in the IFC and OFC states of BbZIP. (**A**) Exposed and buried L200 in the OFC and IFC, respectively. Divalent metal substrates (M^2+^) are depicted as black spheres, and the arrows indicate the movement of the metal substrate in transport. (**B**) Assessment of the accessibility of C200 using thiol reacting agents (NEM and mPEG5K) in the absence and presence of metal substrates Zn^2+^ and Cd^2+^ in Western blot experiments.

### Characterization of the anti-BbZIP monoclonal antibody

In a sandwich ELISA, the coating antibody works in solution to capture the antigen in the native state. Therefore, it is crucial to examine whether the anti-BbZIP antibody is able to bind BbZIP in solution with a high affinity. We purified the anti-BbZIP antibody from hybridoma (**Figure 2A**) and then generated the Fab fragment to test complex formation. The mixture of the purified L200C variant and the Fab fragment was applied to a size-exclusion chromatography and the left-shifted peak, when compared to BbZIP in DDM alone (**Figure 2B**), indicated the formation of a stable BbZIP-Fab complex. Using single-particle cryo-EM, we can see a dumbbell-shaped density protruding on the surface of the DDM micelles (**Figure S1**), consistent with a Fab fragment bound with a detergent solubilized BbZIP. Although we were not able to solve the cryo-EM structure of this Fab-BbZIP complex due to the varied orientations between the two proteins, the result supported a tight binding of the anti-BbZIP antibody to its antigen in the native state, allowing us to use this antibody in the development of the sandwich ELISA.

**Figure 2.**
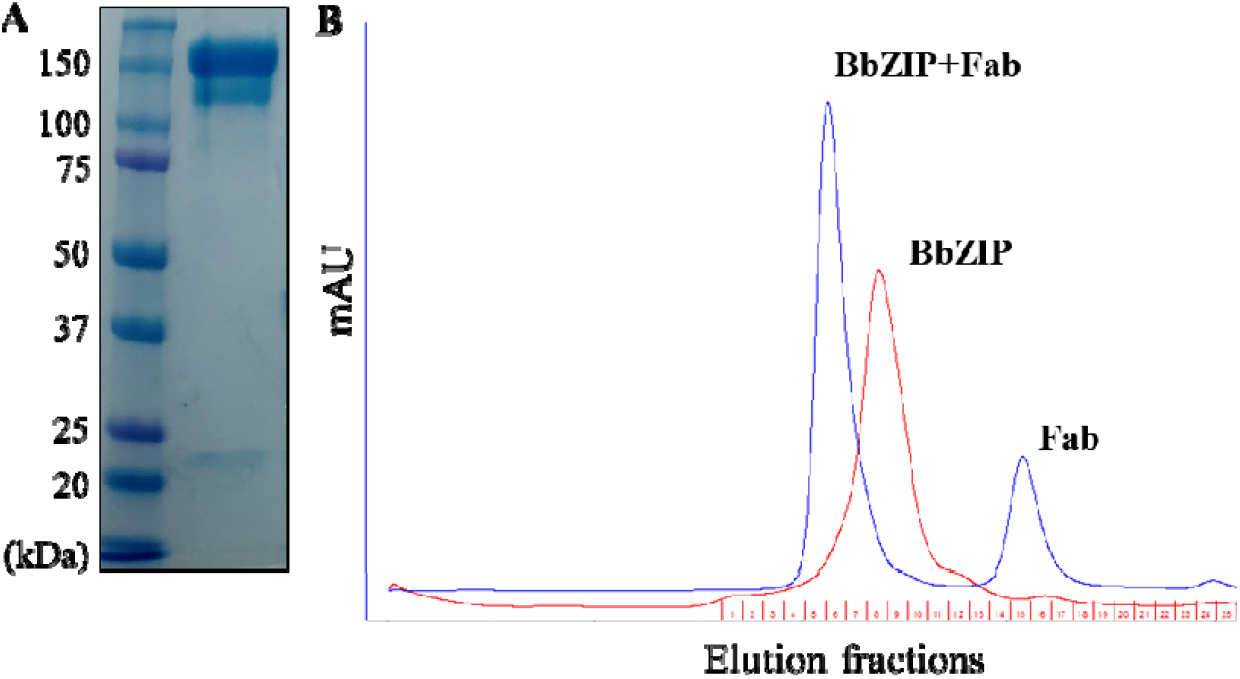
Characterization of the anti-BbZIP monoclonal antibody. (**A**) Non-reducing SDS-PAGE of the purified antibody. (**B**) Stable L200C-Fab complex in solution. The mixture of purified Fab and the L200C variant in DDM was applied to size-exclusion chromatography (blue profile), in comparison with the sample containing only BbZIP (red profile).

### Development of a sandwich ELISA to study metal substrate binding to the transport site

The design of the sandwich ELISA is illustrated in **Figure 3A**. In brief, the membrane fraction of the *E*.*coli* cells expressing the L200C variant is incubated with a biotinylation probe, maleimide-PEG2-biotin, in the absence and presence of metal substrates. This probe can label the protein with a biotin through a selective reaction of the maleimide moiety with exposed cysteine residues, and the hydrophilic (ethylene glycol)_2_ arm between maleimide and biotin allows the latter to be exposed to the protein surface after labeling. Following the labeling reaction, the membrane fraction was washed to remove the residual probe, dissolved in 1% DDM, and added to the 96-well plate pre-coated with the anti-BbZIP antibody to allow the coating antibody to capture the L200C variant. After extensive wash to remove irrelevant proteins from the cell membrane, an HRP-conjugated streptavidin was added to the wells and the later steps till colorimetric analysis using a microplate reader were the same as for the standard ELISA. The membrane fraction of the cells transformed with an empty vector was used as blank. As shown in **Figure 3B**, the L200C sample generated a much stronger signal than the blank sample and the difference between them represented the specific biotin labeling to the single cysteine of the L200C variant. Importantly, the presence of 50 µM Zn^2+^ or Cd^2+^ significantly reduced the signal by approximately 50% (**Figure 3B**), indicating that the labeling efficiency, which reflects the accessibility of C200, was reduced upon the addition of the substrates due to the metal binding induced OFC-IFC switch. This result is consistent with the results of the cysteine accessibility assay studied by using Western blot (**Figure 1**), allowing us to use this rapid and quantitative assay to study the metal substrate binding to the M1 site. Notably, the signals in the presence of the metal substrates were not reduced to the level of the blank, indicating that, although on average C200 is more buried due to the increased population of IFC species in the presence of metal substrates, the OFC species are still present and in an equilibrium with the species in the IFC.

**Figure 3.**
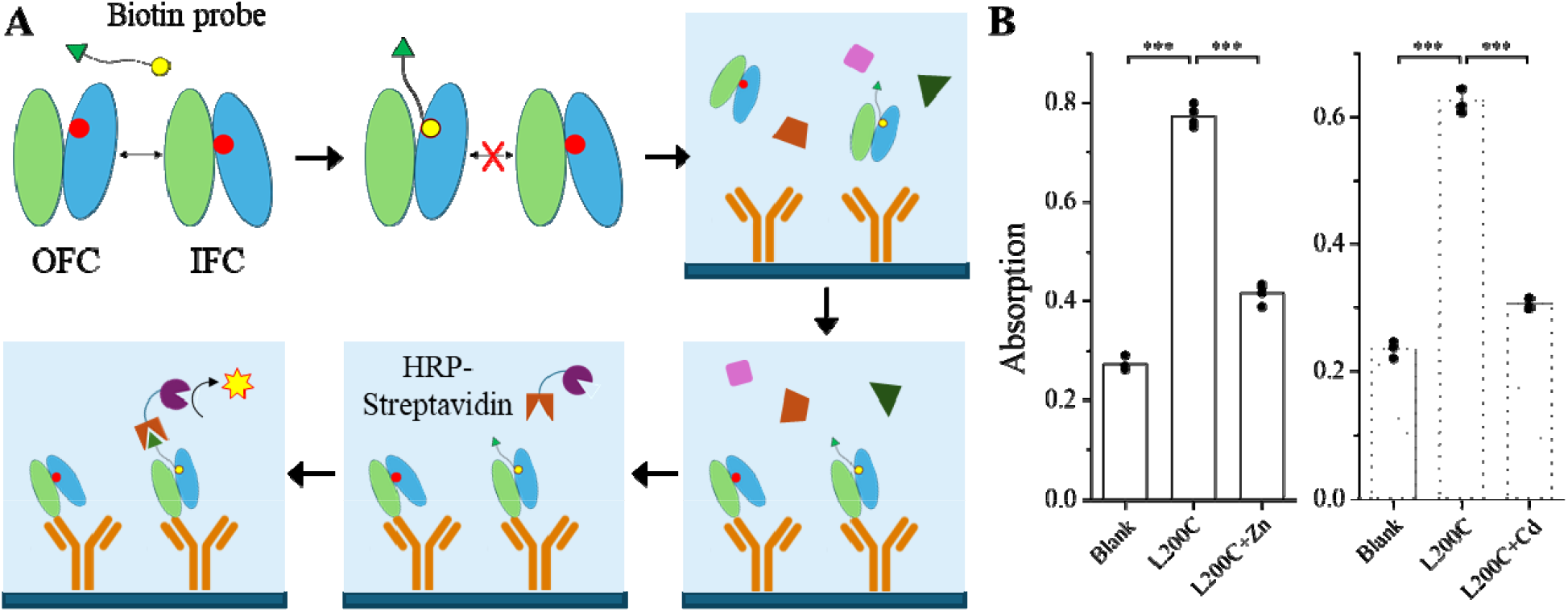
Development of a sandwich ELISA to characterize the conformational state of BbZIP. (**A**) Cartoon illustration of the experimental procedure. The L200C variant of BbZIP in the membrane fraction of *E. coli* cells is incubated with a biotinylation probe, maleimide-PEG2-biotin. Only the protein with the exposed C200 (red dot) is selectively labeled with biotin (green triangle) via maleimide (yellow dot). The DDM solubilized membrane fraction is added to a 96-well plate that have been coated with the anti-BbZIP antibody. After wash, the HRP-conjugated streptavidin is added to selectively bind to the biotin labeled BbZIP that has been caught by the antibody. After the removal of the residual HRP-Streptavidin, the signals generated by the HRP are measured by a plate reader. Adapted from bioRender. (**B**) Detection of the accessibility of C200 in the absence and presence of metal substrates (Zn^2+^ and Cd^2+^ at 50 µM). Each dot represents the result of one out of four replicates that conducted for each condition. ***: *P*<0.001. The *P* values are 1.7×10^−8^, 3.7×10^−7^, 2.8×10^−8^, and 6.5×10^−8^ from left to right.

### Determination of the binding affinities of Zn^2+^ and Cd^2+^ to the M1 site

We then conducted this sandwich ELISA in the presence of different amount of Zn^2+^ and Cd^2+^ in a metal-citrate buffer where the free metal concentrations were calculated using the dissociation constants of the metal-citrate complexes at the website of WEBMAXC (https://somapp.ucdmc.ucdavis.edu/pharmacology/bers/maxchelator/webmaxc/webmaxcS.htm). As shown in **Figure 4A**, the increase of Zn^2+^ reduced the readings in dose-dependent manner for the L200C sample. The readings for the blank under the same condition were also reduced slightly, but the magnitude of the decrease is much smaller than that for the L200C sample. Thus, the reduced signal in the latter can be primarily attributed to the reduced cysteine accessibility upon Zn^2+^ binding. By plotting the signals specifically derived from the L200C variant, which were calculated by subtracting the readings of the blank from those of the L200C samples obtained under the same conditions, against the free concentration of Zn^2+^, a metal binding curve was generated. Curve fitting using a Hill model led to the determination of the apparent dissociation constant for Zn^2+^ (*K*_appa,Zn_=1.9±0.4 µM, mean±S.E., n=3), which in general agrees with the apparent *K*_M_ values of other ZIPs obtained in kinetics studies [11, 12, 17, 18, 34, 42, 43]. The same experiment was also conducted in the presence of Cd^2+^ and the calculated *K*_appa,Cd_ was 0.7±0.1 µM (mean±S.E., n=3) (**Figure 4B**), consistent with the notion that the Cd^2+^-bound BbZIP is more stable than the Zn^2+^-bound state and thus more suitable for structural studies [34]. Indeed, the only Zn^2+^-bound structure of BbZIP was obtained through soaking the Cd^2+^-bound crystals with excess Zn^2+^ [39].

**Figure 4.**
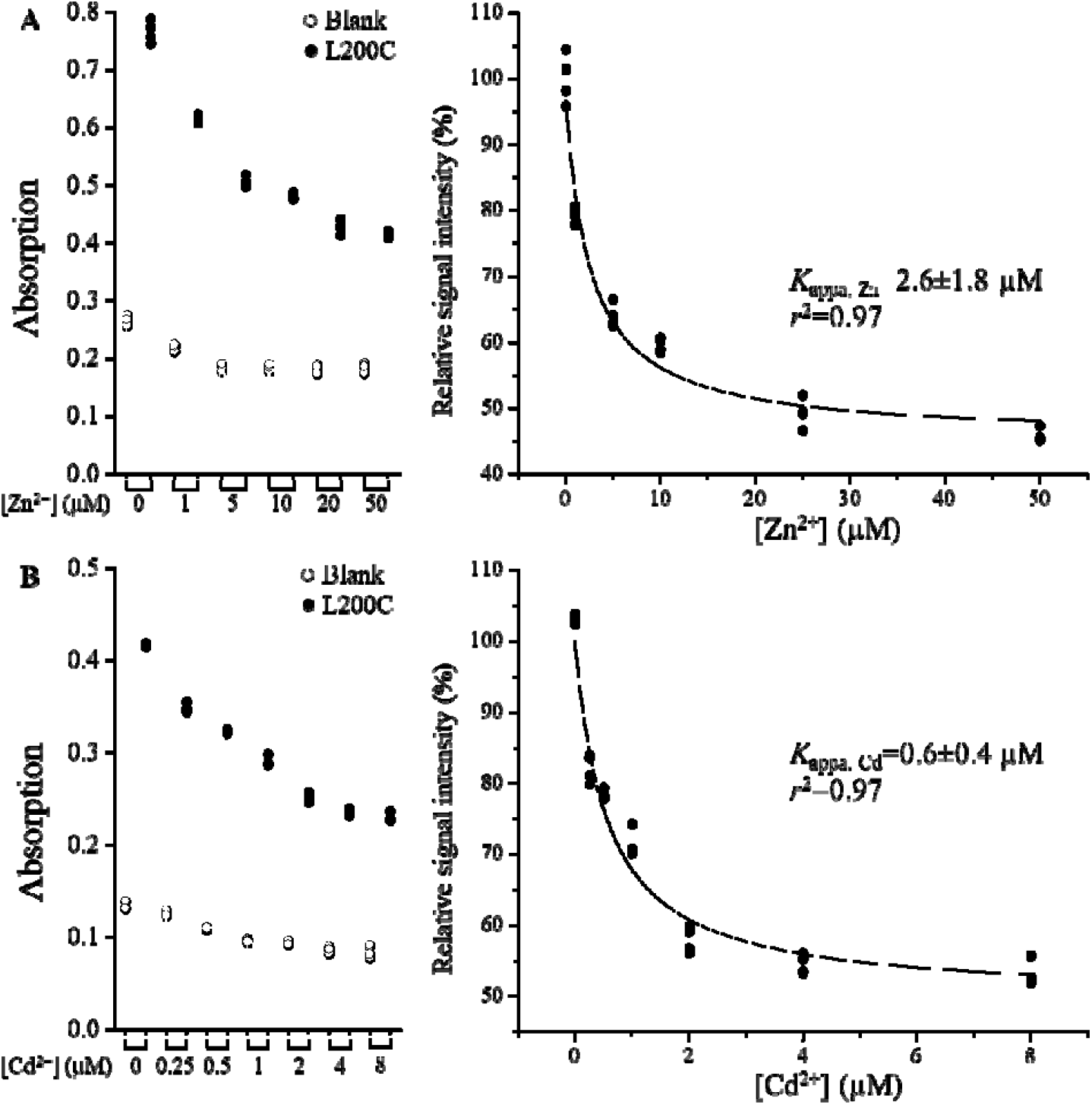
Determination of the dissociation constants of the L200C variant for Zn^2+^ (**A**) and Cd^2+^ (**B**) using the sandwich ELISA. The free M^2+^ concentrations were calculated using the dissociation constants of the metal-citrate complexes at the website of WEBMAXC. The raw data from one experiment with four replicates are shown in the left panel where the readings of the blank (open circle) and the L200C variant (solid circle) were collected in the presence of the same amount of metal substrates. In the right panel, the signals specifically derived from the L200C variant, which were obtained by subtracting the raw data of the blank from those of the L200C variant, were plotted against the free concentration of metal substrates. Curve fitting using a Hill model with n=1 was conducted to estimate the apparent dissociation constant (*K*_appa_) and the results of curve fitting are expressed as best fit ± standard error. *K*_appa, Zn_ and *K*_appa, Cd_ are 1.9±0.4 µM and 0.7±0.1 µM (mean±S.E.), respectively, from three independent experiments.

### Reduced Zn^2+^ binding by a positive charge introduced at the M2 site

With the approach established to study metal binding to the M1 site, we examined the whether a replacement of a negatively charged metal chelating residue at the M2 site with a lysine, which mimics the binding of a divalent metal ion, would have any effect on the binding of Zn^2+^ at the M2 site. As there are two negatively charged residues in the M2 site of BbZIP, D208 and E240 (**Figure 5A**), we introduced two separate mutations into the L200C variant to generate two double variants, L200C/D208K and L200C/E240K, and it seems that the introduced lysine residue can be modeled into the structure without causing clashes (**Figure S2**). Both variants were expressed in the membrane fraction at levels similar to the L200C variant (**Figure 5B**). Using the sandwich ELISA, we found that both variants can be labeled to the levels similar to the L200C variant and addition of Zn^2+^ led to significantly reduced signals (**Figure 5C**), indicating that (1) the M1 site in these variants is similarly exposed to the solvent; and (2) Zn^2+^ can bind to the M1 site and induce a conformational change to reduce the accessibility of C200. However, the calculated *K*_appa_ values showed that the Zn^2+^ binding to the M1 site of both double variants were significantly reduced by 10-20 folds (35±5 µM and 23±3 µM, mean±S.E., n=3, for the L200C/D208K variant and the L200C/E240K variant, respectively) when compared to the L200C variant (1.5±0.1 µM, mean±S.E., n=3) (**Figure 5D**). This drastically diminished Zn^2+^ binding to the M1 site is likely due to the repulsive force between the two positive charges in the transport site, which is consistent with our recent simulation study in which it was shown that a Zn^2+^ at the M2 site facilitates the metal release from the M1 site into the cytoplasm [35]. Accordingly, whether both metal binding sites are used in transport is dependent on the Zn^2+^ concentration in the extracellular/periplasmic space – when the extracellular Zn^2+^ concentration is low, only the M1 site will be used to transport Zn^2+^; whereas when the extracellular Zn^2+^ is elevated, both metal binding sites will be occupied and the metal at the M1 site can be rapidly released into the cytoplasm. However, the physiological significance of this putative two-mode hypothesis remains elusive. As the M2 site of ZupT from *E*.*coli* was reported to preferentially bind and transport Fe^2+^ [45], the two metal binding sites with different preference for metals may provide a mechanism to allow the crosstalk between different metals.

**Figure 5.**
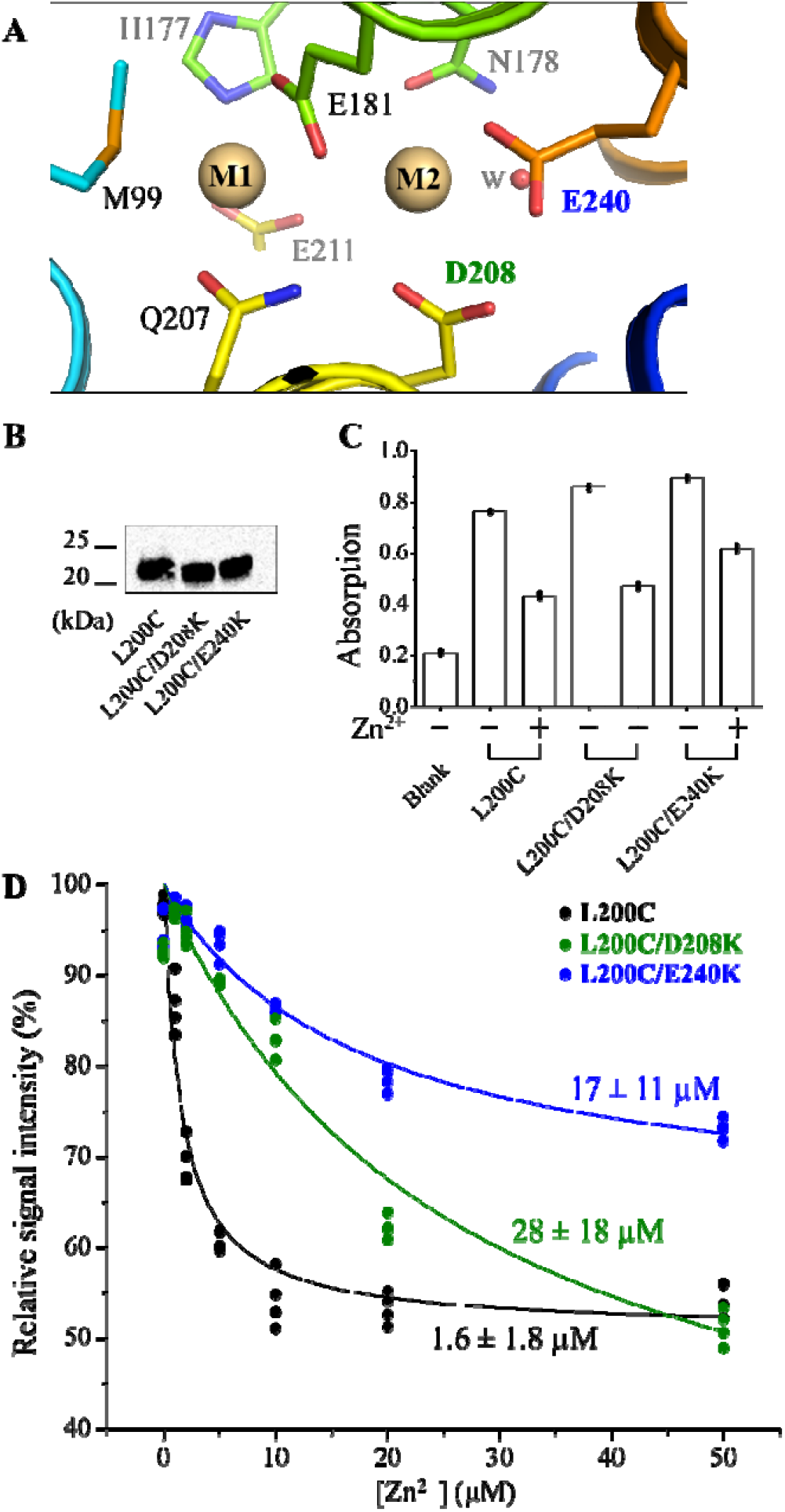
Effects of introducing a lysine residue at the M2 site on the binding of Zn^2+^ to the M1 site of BbZIP. (**A**) The M1 and M2 sites of BbZIP (PDB 5TSB). Cd^2+^ ions are depicted as brown spheres. The metal chelating residues are labeled and shown in stick mode. D208 and E240, the two negatively charged residues in the M2 site are colored differently from other residues. (**B**) Comparison of the expression levels of the variants. The anti-BbZIP was used as primary antibody in the Western blot experiment. (**C**) The representative data of the sandwich ELISA of the three variants. (**D**) Determination of the dissociation constants of the variants for Zn^2+^. The free Zn^2+^ concentration was controlled by using a Zn^2+^-citrate buffer calculated at the website of WEBMAXC. Curve fitting using the Hill model with n=1 was conducted to estimate the apparent dissociation constant (*K*_appa_) and the results of curve fitting are expressed as best fit ± standard error. The *K*_appa, Zn_ values are 1.5±0.1 µM, 35±5 µM, and 23±3 µM (mean±S.E.), for L200C, L200C/D208K, and L200C/E240K, respectively, from three independent experiments.

Another notable observation is that, for the L200C/E240K variant, the signal reduction upon Zn^2+^ addition was smaller than the other two variants (**Figures 5C**&**5D**). Since the signal ratio between the apo and metal-bound states reflects the balance of the IFC-OFC equilibrium, the larger ratio for the L200C/E240K variant (due to a smaller reduction upon Zn^2+^ binding) suggests that this variant favors the OFC more than the other two variants, and thus may be an attractive target for resolving the long-sought structure of the OFC.

### Probing the conformational states of the transport-compromised variants

In our previous study of ZIP4, we showed that replacing small residues at the interface between the transport and scaffold domains with bulky ones resulted in significantly reduced or abolished transport activity [36]. We proposed that this was due to impaired sliding between the two domains when the interface became bumpy, raising the possibility that these variants may be locked into a single conformation and thus be ideal targets for structural biology studies. As these small residues are conserved in many other ZIPs, including BbZIP, we decided to use the sandwich ELISA to examine if any of these variants favors the OFC, as the L200C/E240K variant does (**Figure 5**), so that they could be selected to resolve the OFC structure. Four single mutations were introduced into two positions of the L200C variant, A95 and A203 (**Figure 6A**), resulting in four double variants - A95V/L200C, A95F/L200C, L200C/A203V and L200C/A203F. In ZIP4, the corresponding mutations have been shown to significantly reduce or completely abolish transport activity [36]. All four variants were expressed in the membrane fraction with a similar level of expression to the L200C variant (**Figure 6B**), and they were labelled with more biotin in the absence of Zn^2+^ than in the presence of 50 µM Zn^2+^, indicating that these variants are able to perform the OFC-IFC conformational switch. Calculation of the ratios between the signals with and without Zn^2+^ showed that three variants, including A95F/L200C, A203V/L200C and A203F/L200C, had significantly lower ratios than the L200C variant (**Figure 6C**), suggesting that these variants favor the IFC more than the L200C variant upon Zn^2+^ binding to the M1 site. Although this result does not support further structural characterization of these variants since several structures of the BbZIP in the IFC have been solved [34-39], it demonstrated the value of this assay in rapidly identifying suitable targets for structural biology studies.

**Figure 6.**
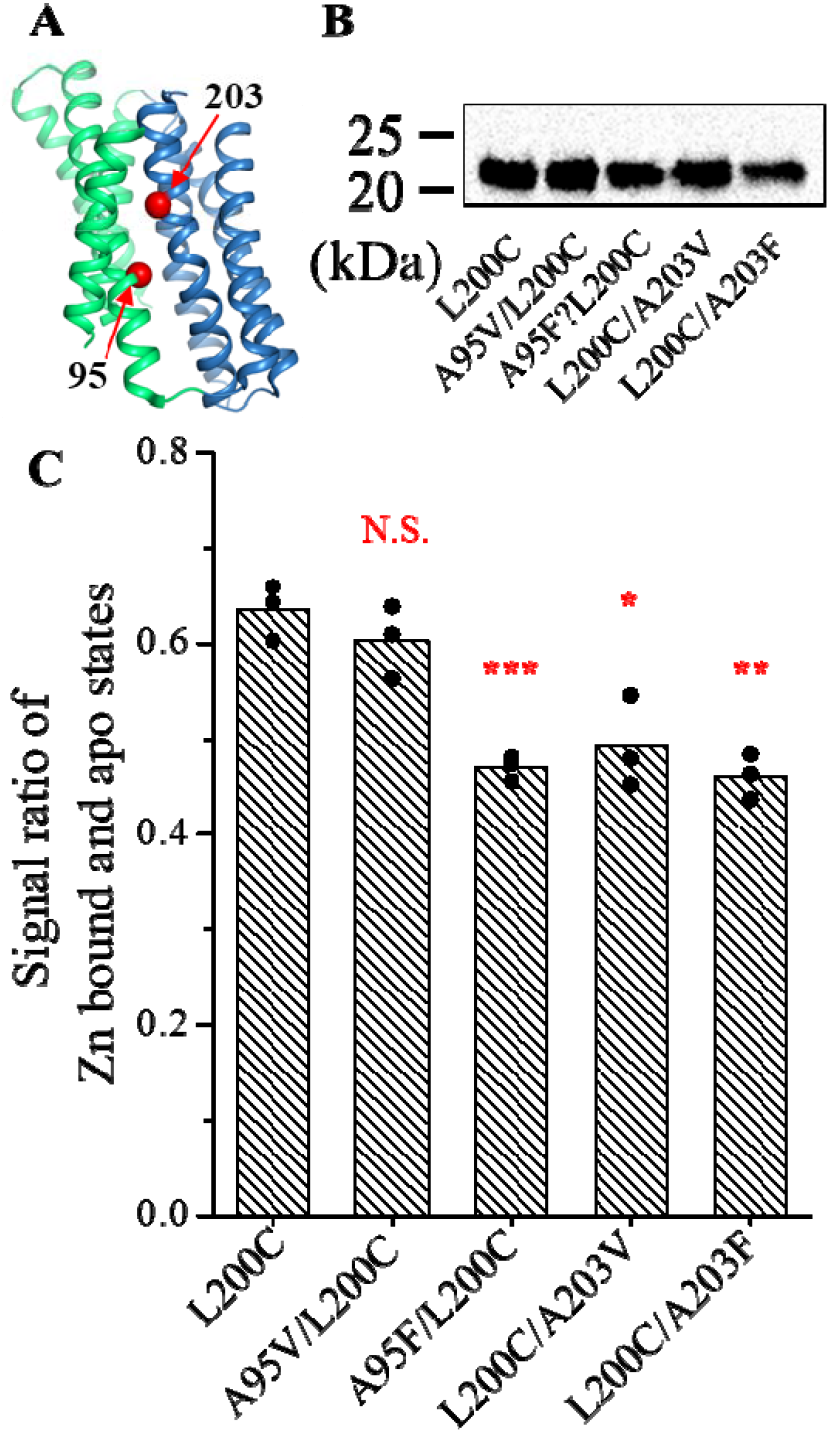
Assessment of the conformational state of the BbZIP variants with perturbed domain interface. (**A**) Location of A95 and A203 in the structure of BbZIP (PDB 5TSB). The Cα atoms of A95 and A203 are shown in red sphere. The scaffold and transport domains are colored in green and blue, respectively. (**B**) Comparison of the expression levels of the variants. The anti-BbZIP was used as primary antibody in the Western blot experiment. (**C**) Ratios of the signals in the presence and absence of 50 µM Zn^2+^ in the sandwich ELISA. The data are from three independent experiments with the data from each experiment shown as one dot. Four replicates were included in one experiment. *: *P*<0.05; **: *P*<0.01; ***: *P*<0.001. The *P* values are 0.32, 0.00084, 0.011, and 0.0013 from left to right in the column chart.

## Conclusion

In this work, we have developed a sandwich ELISA to probe the conformational state of a representative ZIP in the native state. Using this tool, we have reported for the first time the binding affinity of metal substrates, Zn^2+^ and Cd^2+^, to the M1 site of BbZIP; revealed the drastically reduced zinc binding to the M1 site when the neighboring M2 site is occupied by a positive charge, providing new evidence for an auxiliary role of the M2 site in transport; and demonstrated the application of this assay in the identification of targets for subsequent structural study. We anticipate that this approach can be applied to the study of other transporters whose conformational changes are significantly altered by substrate binding.

## Experimental procedures

### Genes, plasmids, mutagenesis, and reagents

The vector harboring the gene encoding BbZIP with an N-terminal His-tag and a thrombin cleavage site in between was the same as reported [34]. Site-directed mutagenesis was performed using the QuikChange® Mutagenesis kit (Agilent), and the primers used for mutagenesis are listed in **Table S1**. NEM, Tris(2-carboxyethyl)phosphine (TCEP), Tween-20, Bovine serum albumin (BSA), phosphate buffered saline (PBS), and other reagents were purchased from Sigma-Aldrich. mPEG5K was purchased from Creative PEGWorks. EZ-Link™ Maleimide-PEG2-Biotin and HRP-conjugated streptavidin (Cata#21130) were purchased from Thermo Fisher Scientific.

### Protein expression and purification

The expression protocol for BbZIP has been previously described [34]. Briefly, the L200C variant were expressed in the C41(DE3)pLysS strain (Lucigen) using LBE-5052 autoinduction medium for 24 hours at room temperature. After harvesting, spheroplasts were prepared and lysed in a buffer containing 20 mM Hepes (pH 7.3), 300 mM NaCl, 0.25 mM CdCl_2_, and cOmplete protease inhibitors (Sigma-Aldrich). To solubilize the membrane fraction, n-Dodecyl-β-D-maltoside (DDM, Anatrace) was added to a final concentration of 1.5% (w/v). His-tagged L200C proteins were purified using HisPur Cobalt Resin (Thermo Fisher Scientific) in a buffer containing 20 mM Hepes (pH 7.3), 300 mM NaCl, 5% glycerol, 0.25 mM CdCl_2_, and 0.1% DDM. The purified protein was concentrated and further purified by size-exclusion chromatography on a Superdex Increase 200 column (GE Healthcare) equilibrated with the buffer containing 10 mM Hepes, pH 7.3, 300 mM NaCl, 5% glycerol, 0.25 mM CdCl_2_, and 0.05% DDM. 1 mM TCEP was included during purification but excluded from size-exclusion chromatography. Peak fractions were pooled for cryo-EM sample preparation.

### Generation of the anti-BbZIP monoclonal antibody and the Fab

The hybridomas secreting the custom anti-BbZIP monoclonal antibody were generated by Creative Biolabs. The antibody was produced in a CELLine bioreactor using the Hybridoma-SFM medium from Thermo Fisher Scientific. After purification from the supernatant of the culture media using Protein G Agarose (Cata#20398, Thermo Fisher Scientific), the antibody was used as the primary antibody in Western blot or processed to generate the Fab.

To prepare the Fab, an immobilized papain resin (Cata#20341, Thermo Fisher Scientific) was incubated with the purification antibody in the digestion buffer containing 100 mM sodium phosphate, pH 7.0, 10 mM EDTA, and 20 mM cysteine with a 1:100 ratio (papain:IgG) at 37 °C for 9 h. The flow through was applied to a Protein A agarose to remove the Fc fragments and the undigested IgG. The antibody can be stored at 4 °C for months without losing the binding capability.

Purified L200C protein was incubated with the Fab at 1:4 mass ratio (L200C:Fab) under gentle shaking at 4□°C overnight and then applied to size-exclusion chromatography on a Superdex Increase 200 column (GE Healthcare) equilibrated with the buffer containing 20□mM Hepes, pH 7.3, 300□mM NaCl, 0.05% DDM. Peak fractions were pooled and concentrated to 2□mg/ml for grid preparation for cryo-EM study.

### Cysteine accessibility assay

Membrane fractions derived from spheroplasts of *E. coli* cells expressing the BbZIP single cysteine variants were incubated with 3 mM NEM for 1 h at 4°C. Excess NEM was removed by washing the samples twice using a buffer containing 100 mM Tris (pH 7.0), 60 mM NaCl, and 10 mM KCl, followed by centrifugation at 13, 000 rpm for 5 min. The membrane fraction was then collected and solubilized in a denaturing buffer containing 6 M urea, 0.5% SDS, and 0.5 mM dithiothreitol to quench residual NEM. This mixture was gently shaken at room temperature for 15 minutes. To label the unmodified cysteine residue, 5 mM mPEG5K was added and incubated for 1 h at room temperature. For control samples without NEM treatment, the membrane fractions were similarly solubilized in the denaturing buffer and treated with 5 mM mPEG5K as described. All samples were mixed with 4x SDS-PAGE sample loading buffer containing 20% β-mercaptoethanol and analyzed by SDS-PAGE. The BbZIP variants were detected by Western blotting using the custom anti-BbZIP monoclonal antibody as the primary antibody and an HRP-conjugated anti-mouse IgG (1:5000 dilution, Cell Signaling Technology, Catalog#7076S) as the secondary antibody. Images of the blots were captured using a Bio-Rad ChemiDoc Imaging System.

### Sandwich ELISA

The membrane fraction from cells expressing the BbZIP variants was incubated in a solution containing 20 mM Hepes, pH 7.3, 100 mM NaCl, and 10 mM sodium citrate for 1 h at 4 °C in the absence or presence of various metals at the indicated concentrations. Citrate was used to form a metal-citrate buffer to control the free concentrations of metal ions in the range of 0-50 µM. Maleimide-PEG2-biotin at 50 µM (final concentration) was added to the membrane fraction to initiate the labeling reaction at 4 °C for 1 h. To remove residual biotin labelling reagent, the membrane fraction was washed four times with the wash buffer containing 20 mM Hepes, pH 7.3, 100 mM NaCl and then dissolved in the wash buffer plus 1% DDM. The solubilized membrane fraction was diluted 10-fold before addition to the 96-well plate for ELISA. The membrane fractions from cells transformed with an empty vector were treated in the same way and used as blank in ELISA.

A 96-well plate was coated overnight at 4°C with 100 µl of anti-BbZIP antibody (10 µg/ml) and washed four times with PBS plus 0.05% Tween-20 (PBST). Wells were blocked with 100 µl of 1% BSA in PBS at 37 °C for 1 hour, followed by four washes with PBST. BbZIP samples were added to each well and incubated at 4 °C for 3 h. After washing, 100 µl of HRP-conjugated streptavidin (at 1:20000 dilution) was added and incubated at 37 °C for 1 h. Wells were extensively washed and 100 µl of 3,3^′^,5,5^′^-tetramethylbenzidine was added and incubated at room temperature for 15–30 min. The reaction was stopped by adding 50 µl of 2 M sulfuric acid, and absorbance was measured at 450 nm using a microplate reader (SpectraMax ABS Plus).

### Statistics

We assumed a normal distribution of the samples and significant differences were examined using Student^’^s t test (two tailed). Uncertainties are reported as S.E., as stated.

## Data availability

All raw and processed data reported in the main text and SI are available upon request. The structure shown in Figures 5, 6, and S1 is retrieved from PDB with the accession code 5TSB.

## Author contributions

Y. Z., R. H., M. S., and J. H. writing–original draft; Y. Z., M. S., and J. H. visualization; Y. Z., M. S., and J. H. validation; Y. Z., R. H., M. S., and J. H. methodology; Y. Z., R. H., and M. S. investigation; Y. Z., R. H., M. S., and J. H. formal analysis; J. H. conceptualization; writing– review and editing; supervision; project administration; funding acquisition.

## Acknowledgments

This work is supported by National Institutes of Health GM140931 (to J. H.). The content is solely the responsibility of the authors and does not necessarily represent the official views of the National Institutes of Health. We thank the youth program “Proteins@MSU” at Michigan State University to recruit Ryan Hu to participate in this work.

## References

1 Krezel, A. and Maret, W. (2016) The biological inorganic chemistry of zinc ions. Arch Biochem Biophys. 611, 3–19

2 Maret, W. (2013) Zinc biochemistry: from a single zinc enzyme to a key element of life. Adv Nutr. 4, 82–91

3 Wessels, I., Fischer, H. J. and Rink, L. (2021) Dietary and Physiological Effects of Zinc on the Immune System. Annual Review of Nutrition. 41, 133–175

4 Laity, J. H., Lee, B. M. and Wright, P. E. (2001) Zinc finger proteins: new insights into structural and functional diversity. Current Opinion in Structural Biology. 11, 39–46

5 Costa, M. I., Sarmento-Ribeiro, A. B. and Gonçalves, A. C. (2023) Zinc: From Biological Functions to Therapeutic Potential. International Journal of Molecular Sciences. 24

6 Kambe, T., Taylor, K. M. and Fu, D. (2021) Zinc transporters and their functional integration in mammalian cells. J Biol Chem. 296, 100320

7 Jeong, J. and Eide, D. J. (2013) The SLC39 family of zinc transporters. Mol Aspects Med. 34, 612–619

8 Eide, D. J. (2020) Transcription factors and transporters in zinc homeostasis: lessons learned from fungi. Crit Rev Biochem Mol Biol. 55, 88–110

9 Hu, J. and Jiang, Y. (2024) Evolution, classification, and mechanisms of transport, activity regulation, and substrate specificity of ZIP metal transporters. Critical Reviews in Biochemistry and Molecular Biology. 59, 245–266

10 Hu, J. (2021) Toward unzipping the ZIP metal transporters: structure, evolution, and implications on drug discovery against cancer. FEBS J. 288, 5805–5825

11 He, L., Girijashanker, K., Dalton, T. P., Reed, J., Li, H., Soleimani, M. and Nebert, D. W. (2006) ZIP8, member of the solute-carrier-39 (SLC39) metal-transporter family: characterization of transporter properties. Mol Pharmacol. 70, 171–180

12 Girijashanker, K., He, L., Soleimani, M., Reed, J. M., Li, H., Liu, Z., Wang, B., Dalton, T. P. and Nebert, D. W. (2008) Slc39a14 gene encodes ZIP14, a metal/bicarbonate symporter: similarities to the ZIP8 transporter. Mol Pharmacol. 73, 1413–1423

13 Liu, Z., Li, H., Soleimani, M., Girijashanker, K., Reed, J. M., He, L., Dalton, T. P. and Nebert, D. W. (2008) Cd2+ versus Zn2+ uptake by the ZIP8 HCO3--dependent symporter: kinetics, electrogenicity and trafficking. Biochem Biophys Res Commun. 365, 814–820

14 Jenkitkasemwong, S., Wang, C. Y., Mackenzie, B. and Knutson, M. D. (2012) Physiologic implications of metal-ion transport by ZIP14 and ZIP8. Biometals. 25, 643–655

15 Winslow, J. W. W., Limesand, K. H. and Zhao, N. (2020) The Functions of ZIP8, ZIP14, and ZnT10 in the Regulation of Systemic Manganese Homeostasis. Int J Mol Sci. 21

16 Korshunova, Y. O., Eide, D., Clark, W. G., Guerinot, M. L. and Pakrasi, H. B. (1999) The IRT1 protein from Arabidopsis thaliana is a metal transporter with a broad substrate range. Plant Mol Biol. 40, 37–44

17 Vert, G., Grotz, N., Dedaldechamp, F., Gaymard, F., Guerinot, M. L., Briat, J. F. and Curie, C. (2002) IRT1, an Arabidopsis transporter essential for iron uptake from the soil and for plant growth. Plant Cell. 14, 1223–1233

18 Nakanishi, H., Ogawa, I., Ishimaru, Y., Mori, S. and Nishizawa, N. K. (2006) Iron deficiency enhances cadmium uptake and translocation mediated by the Fe2+ transporters OsIRT1 and OsIRT2 in rice. Soil Science and Plant Nutrition. 52, 464–469

19 Grass, G., Wong, M. D., Rosen, B. P., Smith, R. L. and Rensing, C. (2002) ZupT is a Zn(II) uptake system in Escherichia coli. J Bacteriol. 184, 864–866

20 Capdevila, D. A., Wang, J. and Giedroc, D. P. (2016) Bacterial Strategies to Maintain Zinc Metallostasis at the Host-Pathogen Interface. Journal of Biological Chemistry. 291, 20858–20868

21 Anzilotti, C., Swan, D. J., Boisson, B., Deobagkar-Lele, M., Oliveira, C., Chabosseau, P., Engelhardt, K. R., Xu, X., Chen, R., Alvarez, L., Berlinguer-Palmini, R., Bull, K. R., Cawthorne, E., Cribbs, A. P., Crockford, T. L., Dang, T. S., Fearn, A., Fenech, E. J., de Jong, S. J., Lagerholm, B. C., Ma, C. S., Sims, D., van den Berg, B., Xu, Y., Cant, A. J., Kleiner, G., Leahy, T. R., de la Morena, M.T., Puck, J. M., Shapiro, R. S., van der Burg, M., Chapman, J. R., Christianson, J. C., Davies, B., McGrath, J. A., Przyborski, S., Santibanez Koref, M., Tangye, S. G., Werner, A., Rutter, G. A., Padilla-Parra, S., Casanova, J. L., Cornall, R. J., Conley, M. E. and Hambleton, S. (2019) An essential role for the Zn(2+) transporter ZIP7 in B cell development. Nat Immunol. 20, 350–361

22 Nolin, E., Gans, S., Llamas, L., Bandyopadhyay, S., Brittain, S. M., Bernasconi-Elias, P., Carter, K. P., Loureiro, J. J., Thomas, J. R., Schirle, M., Yang, Y., Guo, N., Roma, G., Schuierer, S., Beibel, M., Lindeman, A., Sigoillot, F., Chen, A., Xie, K. X., Ho, S., Reece-Hoyes, J., Weihofen, W. A., Tyskiewicz, K., Hoepfner, D., McDonald, R. I., Guthrie, N., Dogra, A., Guo, H., Shao, J., Ding, J., Canham, S. M., Boynton, G., George, E. L., Kang, Z. B., Antczak, C., Porter, J. A., Wallace, O., Tallarico, J. A., Palmer, A. E., Jenkins, J. L., Jain, R. K., Bushell, S. M. and Fryer, C. J. (2019) Discovery of a ZIP7 inhibitor from a Notch pathway screen. Nat Chem Biol. 15, 179–188

23 Li, M., Zhang, Y., Liu, Z., Bharadwaj, U., Wang, H., Wang, X., Zhang, S., Liuzzi, J. P., Chang, S. M., Cousins, R. J., Fisher, W. E., Brunicardi, F. C., Logsdon, C. D., Chen, C. and Yao, Q. (2007) Aberrant expression of zinc transporter ZIP4 (SLC39A4) significantly contributes to human pancreatic cancer pathogenesis and progression. Proc Natl Acad Sci U S A. 104, 18636–18641

24 Weaver, B. P., Zhang, Y., Hiscox, S., Guo, G. L., Apte, U., Taylor, K. M., Sheline, C. T., Wang, L. and Andrews, G. K. (2010) Zip4 (Slc39a4) expression is activated in hepatocellular carcinomas and functions to repress apoptosis, enhance cell cycle and increase migration. PLoS One. 5

25 Zhu, B., Huo, R., Zhi, Q., Zhan, M., Chen, X. and Hua, Z. C. (2021) Increased expression of zinc transporter ZIP4, ZIP11, ZnT1, and ZnT6 predicts poor prognosis in pancreatic cancer. J Trace Elem Med Biol. 65, 126734

26 Taylor, K. M., Vichova, P., Jordan, N., Hiscox, S., Hendley, R. and Nicholson, R. I. (2008) ZIP7-mediated intracellular zinc transport contributes to aberrant growth factor signaling in antihormone-resistant breast cancer Cells. Endocrinology. 149, 4912–4920

27 Hogstrand, C., Kille, P., Nicholson, R. I. and Taylor, K. M. (2009) Zinc transporters and cancer: a potential role for ZIP7 as a hub for tyrosine kinase activation. Trends Mol Med. 15, 101–111

28 Taylor, K. M., Hiscox, S., Nicholson, R. I., Hogstrand, C. and Kille, P. (2012) Protein kinase CK2 triggers cytosolic zinc signaling pathways by phosphorylation of zinc channel ZIP7. Sci Signal. 5, ra11

29 Hogstrand, C., Kille, P., Ackland, M. L., Hiscox, S. and Taylor, K. M. (2013) A mechanism for epithelial-mesenchymal transition and anoikis resistance in breast cancer triggered by zinc channel ZIP6 and STAT3 (signal transducer and activator of transcription 3). Biochem J. 455, 229–237

30 Matsui, C., Takatani-Nakase, T., Hatano, Y., Kawahara, S., Nakase, I. and Takahashi, K. (2017) Zinc and its transporter ZIP6 are key mediators of breast cancer cell survival under high glucose conditions. FEBS Lett. 591, 3348–3359

31 Nimmanon, T., Ziliotto, S., Ogle, O., Burt, A., Gee, J. M. W., Andrews, G. K., Kille, P., Hogstrand, C., Maret, W. and Taylor, K. M. (2020) The ZIP6/ZIP10 heteromer is essential for the zinc-mediated trigger of mitosis. Cell Mol Life Sci

32 Wang, G., Biswas, A. K., Ma, W., Kandpal, M., Coker, C., Grandgenett, P. M., Hollingsworth, M. A., Jain, R., Tanji, K., L□pez-Pintado, S., Borczuk, A., Hebert, D., Jenkitkasemwong, S., Hojyo, S., Davuluri, R. V., Knutson, M. D., Fukada, T. and Acharyya, S. (2018) Metastatic cancers promote cachexia through ZIP14 upregulation in skeletal muscle. Nature Medicine. 24, 770–781

33 Lin, W., Chai, J., Love, J. and Fu, D. (2010) Selective electrodiffusion of zinc ions in a Zrt-, Irt-like protein, ZIPB. J Biol Chem. 285, 39013–39020

34 Zhang, T., Liu, J., Fellner, M., Zhang, C., Sui, D. and Hu, J. (2017) Crystal structures of a ZIP zinc transporter reveal a binuclear metal center in the transport pathway. Sci Adv. 3, e1700344

35 Zhang, Y., Jafari, M., Zhang, T., Sui, D., Sagresti, L., Merz, K. M. and Hu, J. (2024) Molecular insights into substrate translocation in an elevator-type metal transporter. Nature Communications. 15

36 Zhang, Y., Jiang, Y., Gao, K., Sui, D., Yu, P., Su, M., Wei, G. W. and Hu, J. (2023) Structural insights into the elevator-type transport mechanism of a bacterial ZIP metal transporter. Nat Commun. 14, 385

37 Wiuf, A., Steffen, J. H., Becares, E. R., Gronberg, C., Mahato, D. R., Rasmussen, S. G. F., Andersson, M., Croll, T., Gotfryd, K. and Gourdon, P. (2022) The two-domain elevator-type mechanism of zinc-transporting ZIP proteins. Sci Adv. 8, eabn4331

38 Pang, C., Chai, J., Zhu, P., Shanklin, J. and Liu, Q. (2023) Structural mechanism of intracellular autoregulation of zinc uptake in ZIP transporters. Nat Commun. 14, 3404

39 Zhang, T., Sui, D., Zhang, C., Cole, L. and Hu, J. (2020) Asymmetric functions of a binuclear metal center within the transport pathway of a human zinc transporter ZIP4. FASEB J. 34, 237–247

40 Grass, G., Franke, S., Taudte, N., Nies, D. H., Kucharski, L. M., Maguire, M. E. and Rensing, C. (2005) The metal permease ZupT from Escherichia coli is a transporter with a broad substrate spectrum. J Bacteriol. 187, 1604–1611

41 Zhang, T., Sui, D. and Hu, J. (2016) Structural insights of ZIP4 extracellular domain critical for optimal zinc transport. Nat Commun. 7, 11979

42 Zhang, T., Kuliyev, E., Sui, D. and Hu, J. (2019) The histidine-rich loop in the extracellular domain of ZIP4 binds zinc and plays a role in zinc transport. Biochem J. 476, 1791–1803

43 Chowanadisai, W., Graham, D. M., Keen, C. L., Rucker, R. B. and Messerli, M. A. (2013) Neurulation and neurite extension require the zinc transporter ZIP12 (slc39a12). Proc Natl Acad Sci U S A. 110, 9903–9908

44 Pinilla-Tenas, J. J., Sparkman, B. K., Shawki, A., Illing, A. C., Mitchell, C. J., Zhao, N., Liuzzi, J. P., Cousins, R. J., Knutson, M. D. and Mackenzie, B. (2011) Zip14 is a complex broad-scope metal-ion transporter whose functional properties support roles in the cellular uptake of zinc and nontransferrin-bound iron. Am J Physiol Cell Physiol. 301, C862–871

45 Roberts, C. S., Ni, F. and Mitra, B. (2021) The Zinc and Iron Binuclear Transport Center of ZupT, a ZIP Transporter from Escherichia coli. Biochemistry. 60, 3738–3752

