## Supplementary Information for "Probing the substrate binding-induced conformational change of a ZIP metal transporter using a sandwich ELISA"

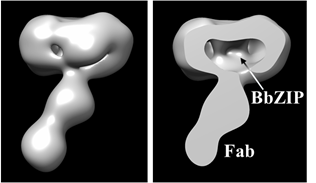

**Figure S1.** Cryo-EM density map of the BbZIP-Fab complex in DDM micelles.

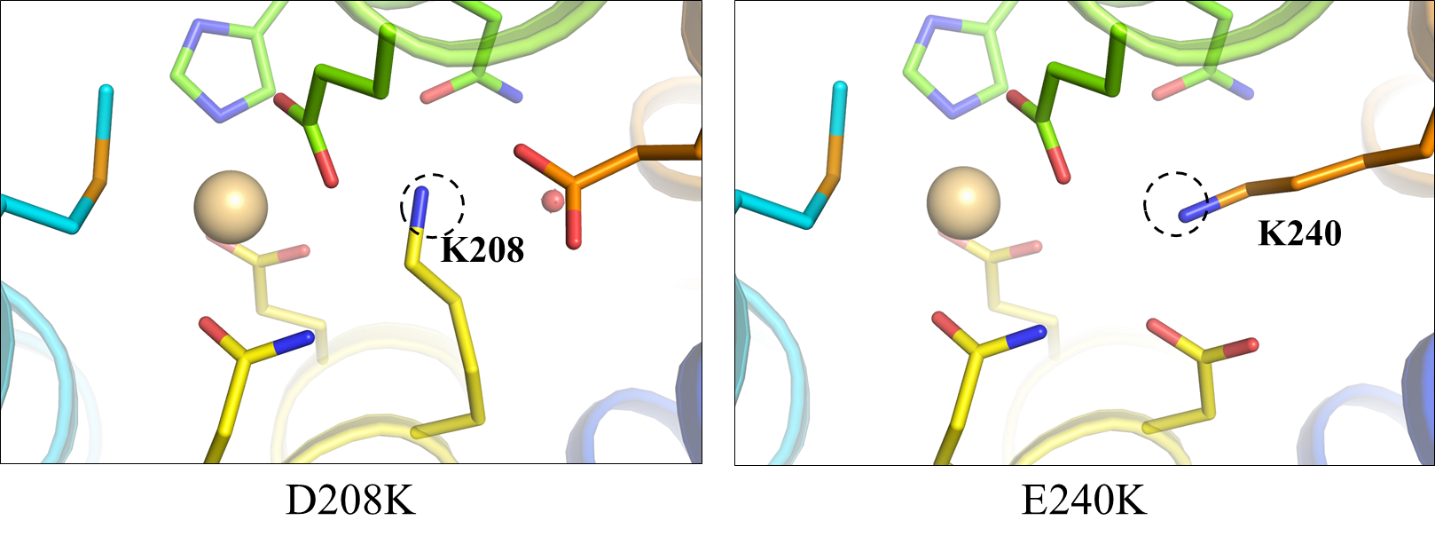

**Figure S2.** Model structures of the BbZIP variants of D208K and E240K. In each variant structure, the lysine residue was manually modelled into the structure of BbZIP to let the primary amine occupy the M2 site (dashed circles) of the Cd-bound structure (PDB 5TSB). No clashes were found when modelling the lysine residues.

**Table S1.** Primers used in this work

| L200C-F | 5’GAT CTG CGT ATT GGT CTG CCG TGC ACC AGC GCC ATT GCA ATT CAG 3’ |
| --- | --- |
| L200C-R | 5’CTG AAT TGC AAT GGC GCT GGT GCA CGG CAG ACC AAT ACG CAG ATC 3’ |
| A95V-F | 5’ CAG GAT GCA ATG CTG GGT TTT GTG GCA GGT ATG ATG CTG GCA GCC3’ |
| A95V-R | 5’ GGC TGC CAG CAT CAT ACC TGC CAC AAA ACC CAG CAT TGC ATC CTG 3’ |
| A95F-F | 5’ CAG GAT GCA ATG CTG GGT TTT TTT GCA GGT ATG ATG CTG GCA GCC3’ |
| A95F-R | 5’GGC TGC CAG CAT CAT ACC TGC AAA AAA ACC CAG CAT TGC ATC CTG3’ |
| A203V-F | 5’ATT GGT CTG CCG CTG ACC AGC GTG ATT GCA ATT CAG GAT GTT CCG 3’ |
| A203V-R | 5’CGG AAC ATC CTG AAT TGC AAT CAC GCT GGT CAG CGG CAG ACC AAT3’ |
| A203F-F | 5’ ATT GGT CTG CCG CTG ACC AGC TTT ATT GCA ATT CAG GAT GTT CCG3’ |
| A203F-R | 5’ CGG AAC ATC CTG AAT TGC AAT AAA GCT GGT CAG CGG CAG ACC AAT3’ |
| N178A-F | 5’ GTT CTG ACC ATT ATC CTG CAT GCG CTG CCG GAA GGT ATG GCA ATT3’ |
| N178A-R | 5’ AAT TGC CAT ACC TTC CGG CAG CGC ATG CAG GAT AAT GGT CAG AAC3’ |
| D208A-F | 5’ACC AGC GCC ATT GCA ATT CAG GCG GTT CCG GAA GGC CTG GCA GTT3’ |
| D208A-R | 5’ AAC TGC CAG GCC TTC CGG AAC CGC CTG AAT TGC AAT GGC GCT GGT3’ |
| E240A-F | 5’ GCC GTT GCA AGC GGT CTG ATG GCG CCG CTG GGT GCC CTG GTT GGT3’ |
| E240A-R | 5’ ACC AAC CAG GGC ACC CAG CGG CGC CAT CAG ACC GCT TGC AAC GGC3’ |
